# Serotonin-2B receptor (5-HT_2B_R) expression and binding in the brain of ageing APP_swe_/PS1_de9_ transgenic mice and in human Alzheimer’s Disease brain tissue

**DOI:** 10.1101/2024.05.30.596651

**Authors:** Marco Anzalone, Sarmad A. Karam, Sanne R. R. Briting, Sussanne Petersen, Majken B. Thomsen, Anne M. Landau, Bente Finsen, Athanasios Metaxas

**Affiliations:** Department of Neurobiology, Institute of Molecular Medicine, University of Southern Denmark, Odense, Denmark; Department of Clinical Research, University of Southern Denmark, BRIDGE - Brain Research - Inter-Disciplinary Guided Excellence, Odense, Denmark; Translational Neuropsychiatry Unit, Department of Clinical Medicine, Aarhus University, Denmark; Department of Life Sciences, School of Sciences, European University Cyprus, Nicosia, Cyprus

**Keywords:** Alzheimer, serotonin, 5-HT_2B_, *APP*_swe_/*PS1*_dE9_, autoradiography, PCR

## Abstract

Although evidence of dysregulation in central serotonergic signaling is widespread in Alzheimer’s disease (AD), relatively little is known about the specific involvement of the serotonin-2B receptor (5-HT_2B_R) subtype. Here, we assessed 5-HT_2B_R expression and binding in brain tissue from *APP*_*swe*_*/PS1*_*dE9*_ transgenic (TG) mice and AD patients. 5-HT_2B_R mRNA was measured by RT-qPCR in 3-to >18-month-old TG and wild-type (WT) littermate mice (n=3-8), and in middle frontal gyrus samples from female, AD and control subjects (n=7-10). The density of 5-HT_2B_Rs was measured by autoradiography using 1 nM [^3^H]RS127445 and 1 μM LY266097. In both mouse and human samples, 5-HT_2B_R mRNA was detectable after 33 amplification cycles. Levels in WT mice, not TG mice, increased with age, and were higher in AD patients compared to control subjects. [^3^H]RS127445 binding was low in the mouse brain, detected after 3 months of age. 5-HT_2B_R density was overall lower in the hippocampus of TG compared to WT mice. Specific binding was too low to be reliably quantified in the human samples. These data provide evidence of a different 5-HT_2B_R expression and binding in the *APP*_*swe*_*/PS1*_*dE9*_ TG model of AD and AD patients. Studies investigating the functional involvement of the 5-HT_2B_R in AD are warranted.

## 1. Introduction

The serotonin-2B receptor (5-HT_2B_R) was the last member of the serotonin receptor-2 family to be sequenced and characterized [1], [2]. It is primarily known for its involvement in peripheral physiological processes, particularly those related to the cardiovascular system and the gastrointestinal tract [1]–[3]. The presence and function of 5-HT_2B_Rs within the central nervous system (CNS), however, have not been thoroughly investigated [4]. Evidence suggests that 5-HT_2B_Rs are present throughout several mammalian brain regions, albeit at relatively low abundance. Their presence has been confirmed in different areas such as the cerebral cortex, dorsal raphe nuclei [5], and amygdala in multiple species, including humans [6]–[8], mice [9], and rats [10]. In addition to neuronal cells, the receptors are known to be expressed by non-neuronal, glial cell populations. Notably, human cortical microglial cells demonstrate a high level of 5-HT_2B_R mRNA expression compared to other cortical cell types [11], [12].

Despite being expressed at low levels in the brain, 5-HT_2B_Rs are considered to play important roles in the regulation of complex emotional behaviors and disease states. For example, the receptors have been implicated in the development of mood and anxiety disorders [13], [14]. In mice, genetically removing or pharmacologically blocking 5-HT_2B_Rs has been reported to increase impulsive behavior and activity, as evidenced in the open field test. Moreover, 5-HT_2B_R knockout mice exhibit schizophrenic negative-, positive- and cognitive-like symptoms, as measured using the social interaction, locomotor response to novelty and novel object recognition tests [14]–[16].

Alzheimer’s disease (AD) is a complex, multifaceted neurodegenerative condition marked by cognitive decline and an array of non-cognitive symptoms [17]. Alterations in serotonergic signaling are a prominent feature of the disorder. Specifically, AD is associated with a pronounced depletion of serotonin levels in several brain regions, including the neocortex and hippocampus [18]–[22]. This serotonergic deficiency is believed to contribute to the cognitive impairment and mood disturbance that are characteristic of the disease. Furthermore, *post-mortem* examinations of brains from AD patients, as well as of animal models of the condition, have demonstrated changes in the expression patterns and/or reductions in the expression of several serotonin receptors [23], [24], including the 5-HT_1A_ [25], 5-HT_2A_ [26]–[29], and 5-HT_6_ receptors [29] and the serotonin transporter [30], [31].

Given the diverse cell types expressing 5-HT_2B_Rs and their extensive and rather under-characterized distribution within the brain, it is plausible that these receptors may influence a range of pathological features in AD, from neuroinflammatory responses to neuronal integrity and function. Here, we employed quantitative polymerase chain reaction (qPCR) and autoradiography to investigate age-related changes in the mRNA expression and binding levels of 5-HT_2B_Rs within the *APP*_*swe*_*/PS1*_*dE9*_ mouse model of AD and in brain tissue from AD patients. Examining the regulation of 5-HT_2B_Rs in AD may shed light on the serotonergic mechanisms that contribute to disease progression.

## 2. Results

### 2.1 5-HT_2B_R mRNA levels in mouse and human brain sections

The mRNA levels of the 5-HT_2B_R were measured in brain sections from age-matched *APP*_*swe*_*/PS1*_*dE9*_ TG and WT animals and in frontal cortex tissue (Brodmann area 46) obtained from confirmed AD patients and control subjects. Alzheimer’s amyloid plaque pathology was confirmed by immunohistochemistry for amyloid-b (Aβ) peptides, using the 6E10 antibody (Figure 1). PCR products of undiluted cDNA from mouse and human samples were determined after 33.1±0.1 and 33.8±0.4 cycles, respectively. In all cases, a single peak was obtained by melt-curve analysis, and no signal was detected in the genomic DNA and buffer controls. In mice (Figure 2A), there was a significant age x genotype interaction effect [F_(3, 29)_ = 5.6; *P*=0.003] on the brain mRNA levels of 5-HT_2B_Rs, which were two-three-fold lower in TG vs. WT mice at >18 months of age (*P*=0.02; Bonferroni post-hoc tests). No main effects of age [F_(3, 29)_ = 1.28; *P*=0.29] and genotype were detected by two-way ANOVA [F_(1, 29)_ = 0.03; *P*=0.95]. In WT mice only, one-way ANOVA revealed an effect of age on the expression level of 5-HT_2B_R mRNA [F_(3, 15)_ = 5.69; *P*=0.01], which was two-three-fold higher in 18-month-old WT mice vs. 3- (*P*=0.03) and 6-month-old animals (*P*=0.03; Bonferroni post-hoc tests). In human samples (Figure 2B), 5-HT_2B_R mRNA was significantly elevated in AD compared to controls subjects [t_(14)_=2.60, *P*=0.02; two-tailed t-test].

**Figure 1.**
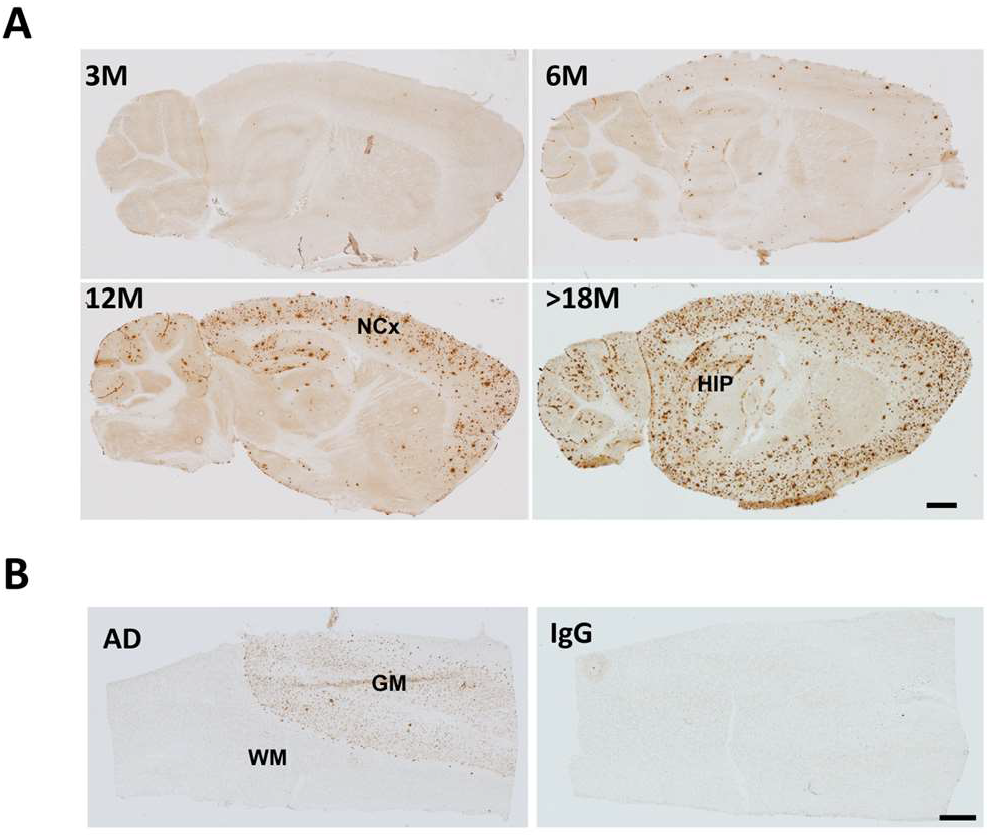
Representative images of Aβ pathology in (A) ageing *APP*_*swe*_*/PS1*_*dE9*_ mice and (B) human AD subjects. Abbreviations: NCx: Neocortex; HIP: Hippocampus; GM: Gray matter; WM: White matter. Scale bars: 1.5 mm (A), 2 mm (B).

**Figure 2.**
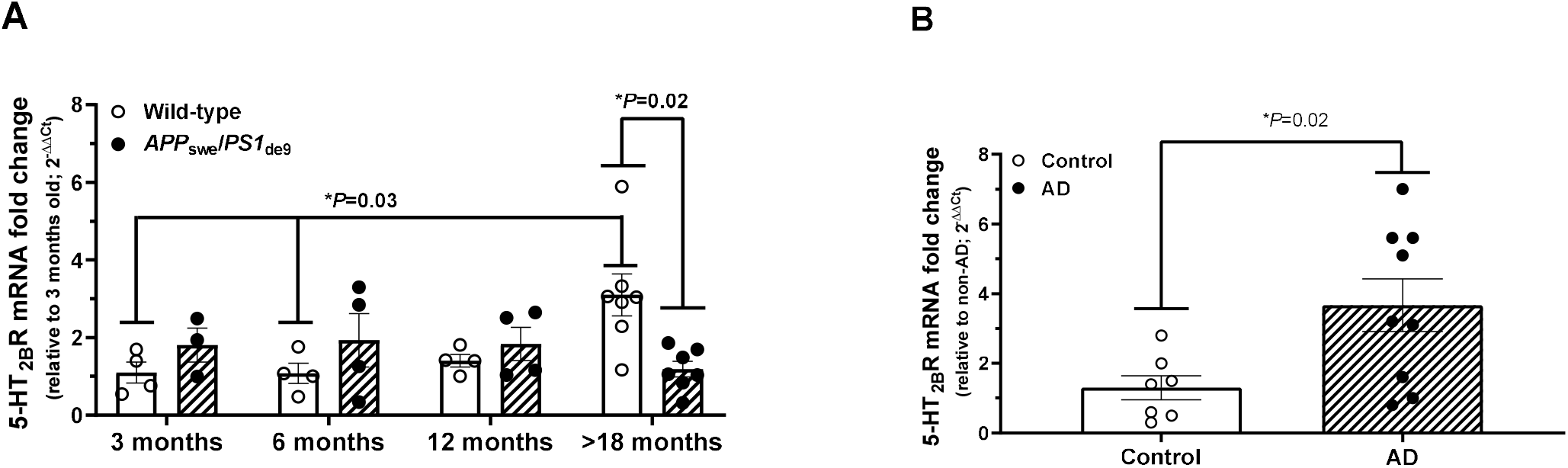
5-HT_2B_R mRNA expression levels in ageing *APP*_*swe*_*/PS1*_*dE9*_ transgenic mice and in human AD subjects. (A) The expression of 5-HT_2B_R mRNA was lower in transgenic (TG) vs. wild-type (WT) mice at >18 months of age. (B) The mRNA expression of 5-HT_2B_Rs was significantly elevated in the AD group compared to the control group. All data are presented as mean ± SEM and normalized to the HPRT-1 reference gene and to the values of 3-month-old WT mice (A), and human control samples (B). Each circle on the plots represents a single data point. Statistical analysis was performed using GraphPad Prism software (ver. 8.2).

### 2.2 [^3^H]RS-127445 autoradiography

The 5-HT_2B_R selective ligands [^3^H]RS-127445 and LY-266097 were used to evaluate the density of 5-HT_2B_Rs in sections obtained from TG and WT mice, AD subjects and non-demented controls. In WT mice, specific [^3^H]RS-127445 binding amounted to ∼15-25% of total binding values in both heart and brain tissue. Levels of specific binding were up to 4-fold higher in the mouse heart, included as a positive control, compared to mouse brain. In human brain sections, levels of specific binding in the gray and white matter were 2.9% and 5.4%, respectively, of total binding, precluding the possibility of accurately quantifying the density of 5-HT_2B_Rs (Figure 3 A-C).

**Figure 3.**
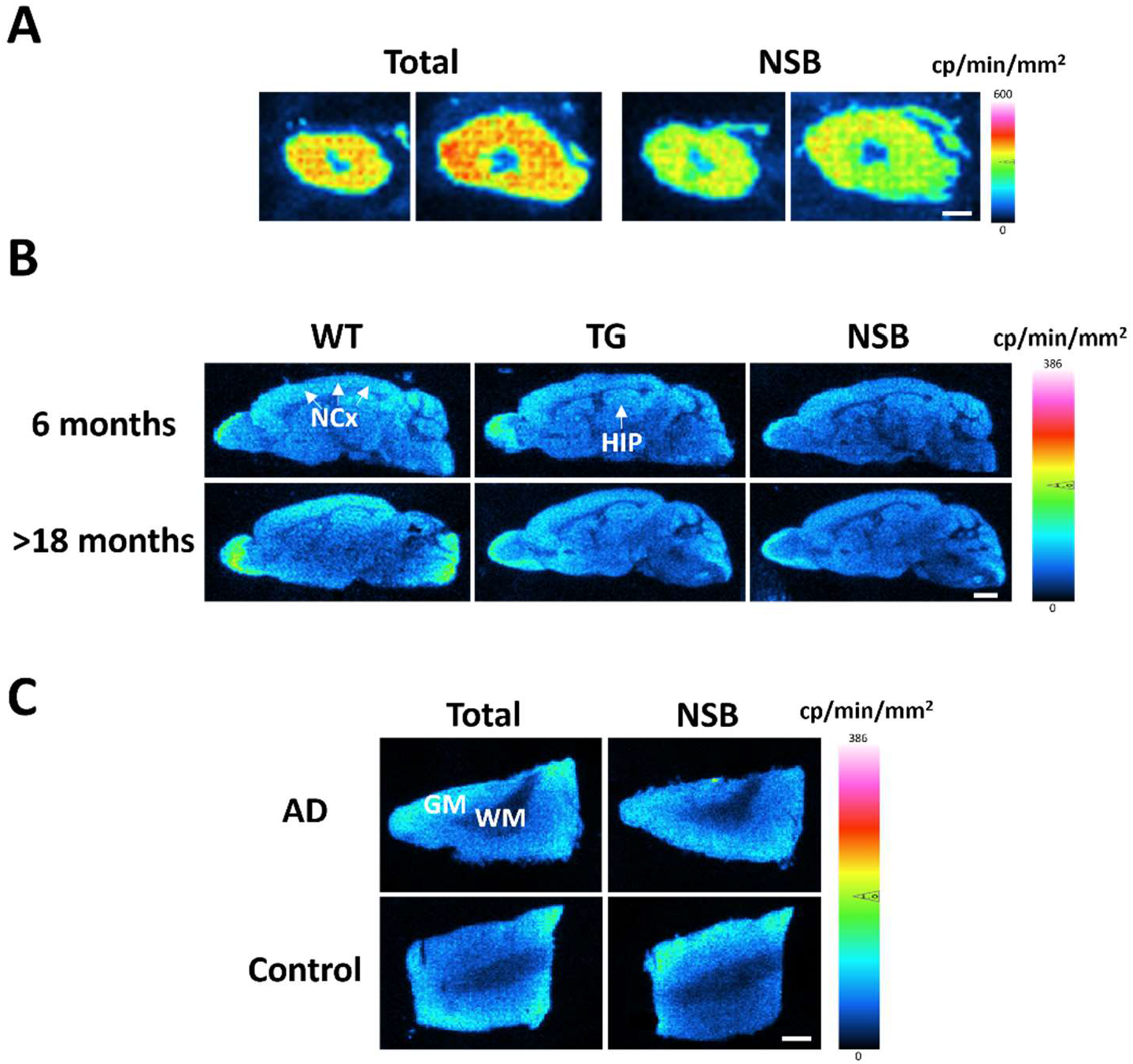
Representative autoradiograms of 1 nM [^3^H]RS127445 binding to (A) mouse heart and (B) mouse brain tissue, and (C) frontal cortex from AD and control cases. Specific binding was defined in the presence of the selective 5-HT_2B_R ligand LY-266097 and amounted to ∼15-25% of total binding values in mouse tissue, and <3% of total binding values in human tissue. Note the high expression of 5HT_2B_R in mouse heart compared to brain tissue. Abbreviations: NCx: Neocortex; HIP: Hippocampus; GM: Gray matter; WM: White matter; NSB: Non-specific binding. Scale bars: 1.25 mm (A), 1.5 mm (B), and 2 mm (C).

No specific binding was detected in the neocortex and the hippocampus of 3-month-old WT and TG animals. In 6-, 12- and >18-month-old mice, specific [^3^H]RS-127445 binding was rather low (<5 cpm/mm^2^). There was no effect of age [F_(2, 25)_ = 0.59; *P*=0.56] and genotype [F_(1, 25)_ = 2.0; *P*=0.17], and no genotype x age interaction effect [F_(2, 25)_ = 0.11; *P*=0.90] on the binding levels of [^3^H]RS-127445 in the neocortex (Figure 4A). In the hippocampus, the binding of [^3^H]RS-127445 was overall lower in TG vs. WT mice. Two way-ANOVA showed an effect of genotype on the density of 5-HT_2B_Rs [F_(1, 25)_ = 5.60; *P*=0.03], with no age [F_(2, 25)_ = 0.065; *P*=0.94] or age x genotype interaction effects [F_(2, 25)_ = 0.49; *P*=0.62]. In line with the *post-hoc* comparisons, showing no differences in the density of 5-HT_2B_Rs between TG and WT mice (Figure 4B), the genotype effect was small.

**Figure 4.**
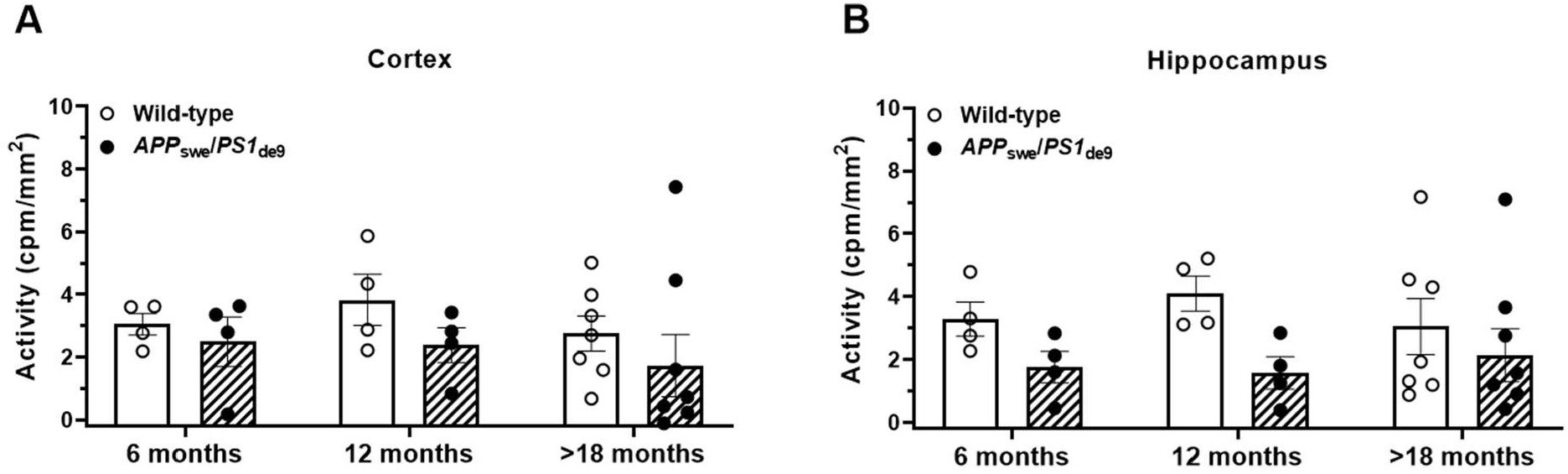
Digital real-time autoradiography of 5-HT_2B_R binding sites in mouse brain tissue. Quantification of [^3^H]RS127445 binding in the neocortex and the hippocampus of ageing WT and *APP*_*swe*_*/PS1*_*dE9*_ TG mice. Values represent the mean specific binding of [^3^H]RS127445 ± SEM in 4-8 animals/group. For each animal, specific binding was determined in five-six consecutive sections. There was an overall effect of genotype on the binding levels of [^3^H]RS127445 in the hippocampus, which were lower in *APP*_*swe*_*/PS1*_*dE9*_ TG vs. WT mice. Specific binding was not detected in 3-month-old mice.

In the cerebellum, low levels of specific binding were first observed by 3 months of age (Supplementary Figure S1). There were no effects of age [F_(3, 29)_ = 1.71; *P*=0.19] and genotype [F_(1, 29)_ = 2.36; *P*=0.14], and no genotype x region interaction effects [F_(3, 29)_ = 1.20; *P*=0.33] on the binding levels of [^3^H]RS-127445 in the cerebellum.

## 3. Discussion

This study was designed to examine the expression of the 5-HT_2B_R subtype within the context of AD pathophysiology. By using the *APP*_*swe*_*/PS1*_*dE9*_ mouse model of AD and human tissue and employing autoradiography and PCR analyses, we present evidence that 5-HT_2B_R mRNA expression is regulated by normal ageing and 5-HT_2B_R binding by Ab plaque pathology. In addition, we report on elevated expression of 5-HT_2B_R mRNA in human AD compared to control subjects.

Early studies on the brain expression levels of the 5-HT_2B_R reported discrepant findings. While some studies documented the presence of 5-HT_2B_R mRNA in mice [9], [32], [33] and rats [3], [5], [34], others failed to detect 5HT_2B_R expression in either species [35]–[38]. To date, the consensus in the scientific literature is that the 5-HT_2B_R mRNA and protein is expressed in the brain of several species, including humans [2], [3], [39], monkeys, cats [37] and dogs [40], albeit at low levels compared to the periphery. The results of our study are consistent with this literature, as they demonstrate low levels of mRNA expression in the human frontal cortex and the brain of C57BL/6J mice.

At the protein level, detection of the 5-HT_2B_R in brain tissue has typically relied on immunoreactivity assays, with studies reporting distribution of the 5-HT_2B_R in the rat and mouse cerebellum, as well as in the rat amygdala, hypothalamus, lateral septum and the frontal cortex [10], [33]. Our study confirms and extends these findings, demonstrating the presence of 5-HT_2B_R protein in the murine neocortex, hippocampus and cerebellum by means of quantitative autoradiography, using the 5-HT_2B_R-selective ligand [^3^H]RS127445. The binding levels of 5-HT_2B_Rs in the mouse brain are low, which aligns well with previously reported low levels of mRNA expression in rats and mice, as well as with our PCR data. Of note, 5-HT_2B_R binding was almost four-fold lower in the mouse brain compared to cardiac tissue, which aligns with the known distribution of the 5-HT_2B_R in the CNS compared to the periphery.

Our study further reveals an age-related modulation of 5-HT_2B_R expression and binding in WT mice, with reliable detection of receptor protein in the rodent neocortex and hippocampus (but not the cerebellum) manifesting only after three months of age. This age-dependent, and perhaps region-specific, regulation might indicate a developmental or physiological adaptation that could influence 5-HT_2B_R availability during a rodent’s lifetime. While the literature on the age-dependent regulation of 5-HT_2B_Rs is sparse, one study reported age-induced increases in the mRNA and protein levels of 5-HT_2B_Rs in 3-30-month-old wild-type mice [41], an observation in line with our PCR analyses and autoradiography results in WT mice. The purported age-dependency in the regulation of 5-HT_2B_Rs might contribute to the inconsistencies in detecting these receptors across different studies within the same species, as previously discussed. Notably, the literature identifies microglia as a major source of 5-HT_2B_Rs in both murine and human neocortex [11], [12], [42], [43]. Additionally, microglia exhibits a several-fold increase in 5-HT_2B_R mRNA levels in response to focal cerebral ischemia [42], compared to other brain cell types, including astrocytes. These findings underscore that microglial 5-HT_2B_R mRNA expression can be regulated by pathological stimuli. Therefore, it is intriguing to speculate that alterations in the mRNA and protein levels of 5-HT_2B_Rs in the cerebral cortex of ageing mice might mirror microglial activation state, both in the context of physiological ageing and AD.

Despite the detection of 5-HT_2B_R mRNA in both mouse and human cerebral cortex, the quantification of specific 5-HT_2B_R binding in the human sections proved difficult. This difficulty may primarily arise from the low levels of 5-HT_2B_R [44]. Technical challenges may stem from the longer *post-mortem* interval and the lower overall quality of the human-derived tissue, which may undermine the integrity of the 5-HT_2B_Rs and the reliability of the binding assay. Moreover, despite the similarities between mouse and human 5-HT_2B_Rs, potential species-specific differences in receptor expression, and pharmacology/binding properties, may have prevented the successful application of [^3^H]RS127445 autoradiography to the human samples [8], [45]. In addition, the density of 5-HT_2B_Rs might be lower in human than in mouse brain sections, which would require optimised protocols to discern specific binding signals from background noise levels. It is thus likely that several different factors have contributed to the difficulties we encountered in quantifying specific 5-HT_2B_R binding in the human neocortex.

The pathophysiology of AD is marked by several important alterations in the serotonergic system. AD and mouse models of Ab plaque pathology are both characterized by loss of serotonergic neurons [46], decreased serotonin synthesis and release [18], serotonin transporter dysfunction [31], as well as altered 5-HT receptor expression and function [23], [24]. While research into the involvement of 5-HT in AD is extensive, the specific role of the 5-HT_2B_R subtype is unknown. Here, we show that the expression of the 5-HT_2B_R mRNA is lower in the brain of aged, >18-month-old TG compared to WT mice, indicating that the effect of Ab plaque pathology on the serotonergic system extends to involve 5HT_2B_R mRNA or the cells expressing the 5HT_2B_R mRNA, which for both mouse and human neocortex are mainly the microglial cells [11], [12], [47]. Additionally, at the protein level, we observed a subtle yet significant overall decrease in the binding levels of 5-HT_2B_R in the hippocampus of TG mice, a finding that further implicates this receptor subtype in Ab pathology, although this trend was observed also at younger ages.

It is noteworthy that PCR analysis of the 5HT_2B_R mRNA in the human frontal cortex showed elevated, rather than lower levels of these receptors in AD. However, the histopathology in Braak stage VI patients is considerably more complex compared to >18-month-old *APP*_*swe*_*/PS1*_*dE9*_ TG mice, which may explain the discrepancy. The elevated 5HT_2B_R mRNA expression in the AD subjects may besides Ab plaque pathology, reflect the pronounced tau pathology and neurodegeneration among other leading to cortical atrophy, which is not observed in the *APP*_*swe*_*/PS1*_*dE9*_ TG mice [48], [49]. Another plausible explanation for the observed differences in the expression of 5-HT_2B_R mRNA between AD subjects and aged TG mice could be interspecies variations in the function and characteristics of microglia, which are major expressors of 5-HT_2B_Rs in both species [11], [12], [43]. It is conceivable that the density, morphology and activation state of the microglial cells may differ significantly between AD subjects and TG mice, contingent upon the disease state. Such disparities in microglial functional characteristics could contribute to the observed differences in 5HT_2B_R mRNA expression between the two species. Clearly, further research is needed to elucidate the regulation of 5HT_2B_R mRNA expression and 5HT_2B_R binding observed in this study.

One limitation of the present investigation is the relatively small sample size, emphasizing the importance of validating our findings in a larger cohort, to increase statistical robustness and enhance the generalizability of our results. The low levels of specific binding in the autoradiography assay present another significant challenge, making it difficult to distinguish the signal of interest from background noise, and thus limiting the sensitivity of our assay to quantify receptors with low expression levels, like the 5-HT_2B_R. Despite these limitations, the specificity of the 5-HT_2B_R signal detected in this study is substantiated by the instrument’s sensitivity threshold and the utilization of two chemically distinct and highly selective ligands to define specific binding, thereby reinforcing the validity of our observations.

In conclusion, this study provides evidence of dysregulated 5-HT_2B_R expression and binding in AD. Given that recent data implicate 5-HT_2B_Rs in modulating neurotransmitter release, synaptic plasticity and neuroinflammation [50]–[54] – processes dysregulated in AD – it is possible that these receptors are involved in the pathogenesis and progression of the disorder. Further investigation into the regulation and function of the 5-HT_2B_R in *APP*_*swe*_*/PS1*_*dE9*_ TG mice and in AD is warranted.

## 4. Materials and Methods

### 4.1 Animals and housing conditions

*APP*_swe_/*PS1*_dE9_ transgenic (TG) [55] and wild-type (WT) littermate control mice were used at 3, 6, 12 and >18 months of age (n=3-8/group; total number of mice: 38; male: 2, female: 36). The mice were bred and maintained in the Biomedical Laboratory of the University of Southern Denmark, on a C57BL/6J background. The animals were group-housed with same-sex siblings within a controlled environment, at a temperature of 21 ± 1 °C and humidity between 45% and 65%. The mice adhered to a 12/12-hour light/dark cycle, with lights on at 7 am, and were provided access to food and water *ad libitum*. Animals were euthanised by cervical dislocation and the brains were promptly extracted, placed on petri dish filled with ice and longitudinally bisected. The right hemisphere was immediately frozen by immersion in isopentane on dry ice (−30°C) and kept at -80°C until used in autoradiography and PCR experiments. The hearts from 2-3 control mice were also obtained and used in autoradiography for positive control purposes.

### 4.2 Human brain sections

To study the binding and expression of 5-HT_2B_R in human brain sections, frozen samples from the middle frontal gyrus (Brodmann area 46) of female AD patients (n=10) and sex- and age-matched non-demented controls (n=7) were obtained from the Netherlands Brain Bank (NBB), Amsterdam, the Netherlands. All AD samples met the criteria outlined in the Braak staging for a definitive diagnosis of AD. Subject characteristics are shown in Table 1.

**Table 1.** Subject characteristics. Braak’s scheme typically consists of six stages, ranging from Stage I (minimal pathology) to Stage VI (severe pathology), which correspond to progressively advanced levels of Alzheimer’s disease tau pathology in the brain. PMI: Post-mortem interval.

| No | Age (years) | PMI (h) | Braak Stage | Study Group |
| --- | --- | --- | --- | --- |
| 1 | 57 | 5.1 | 0 | Control |
| 2 | 94 | 4.1 | I |  |
| 3 | 51 | 5.4 | I |  |
| 4 | 79 | 10.3 | I |  |
| 5 | 75 | 5.3 | I |  |
| 6 | 86 | 7.1 | I |  |
| 7 | 59 | 7.1 | 0 |  |
| **Mean ± SEM** | **71.6±6.1** | **6.3±0.8** | **Median Braak Stage: I** |  |
| 8 | 84 | 7.2 | V | AD |
| 9 | 82 | 4.4 | V |  |
| 10 | 84 | 4.1 | VI |  |
| 11 | 74 | 5.3 | V |  |
| 12 | 82 | 5.3 | VI |  |
| 13 | 73 | 5.3 | VI |  |
| 14 | 80 | 4.2 | VI |  |
| 15 | 75 | 5.0 | V |  |
| 16 | 77 | 6.1 | VI |  |
| 17 | 84 | 4.5 | VI |  |
| **Mean ± SEM** | **79.5±1.4** | **5.1±0.3** | **Median Braak Stage: VI** |  |

### 4.3 Aβ immunohistochemistry

The biotinylated 6E10 antibody (SIG-39340, Nordic BioSite) was used to investigate Aβ pathology in TG mouse and human AD tissue. Clone 6E10 is raised against amino acids 1-16 of human Aβ, recognizing multiple Aβ peptides and precursor forms (manufacturer information). The sections were immersed in 70% formic acid for 30 min, rinsed for 10 min in 50 mM Tris-buffered saline (TBS, pH 7.4), and further washed/permeabilised in TBS containing 1% Triton X-100 (3 x 15 min). Sections were subsequently blocked for 30 min in TBS, containing 10% fetal bovine serum (FBS). Incubation with the 6E10 anti-β-amyloid antibody was carried out overnight at 4°C, in TBS containing 10% FBS (1:500 dilution of stock). Adjacent sections were incubated with biotin-labelled mouse isotype (IgG1) control (MG115, Thermo Fisher Scientific), diluted to the same protein concentration as the primary 6E10 antibody. Following incubation with 6E10, the sections were adjusted to room temperature (RT) for 30 min and washed in TBS+1% Triton X-100 (3 x 15 min). Endogenous peroxidase activity was quenched for 20 min in a solution of TBS/methanol/H_2_O_2_ (8:1:1). After washing in TBS+1% Triton X-100 (3 x 15 min), sections were incubated for 3 h with HRP-streptavidin in TBS/10% FBS (1:200; GE Healthcare Life Sciences). After a final wash in TBS (3 x 10 min), peroxidase activity was visualised with 0.05% 3,3′diaminobenzidine (DAB) in TBS buffer, containing 0.01% H_2_O_2_ (Sigma Aldrich Co.). The developed sections were thoroughly washed in dH_2_O, dehydrated in graded alcohols, cleared in xylene, and cover-slipped with PERTEX^®^ (Histolab Products AB).

### 4.4 5-HT_2B_R mRNA real-time-qPCR

RNA was isolated from human, WT and TG mouse brain sections using Trizol®. Two μg RNA were converted to cDNA using the Applied BiosystemsTM high-capacity cDNA transcription kit (Thermo Fisher Scientific). Triplicate 20 μL samples were subjected to analysis on a StepOne-PlusTM Real-Time PCR system (Applied BiosystemsTM, Thermo Fisher Scientific). Each sample included nuclease-free H_2_O (Thermo Fisher Scientific), 1× Maxima SYBR™ green/probe master mix (Thermo Fisher Scientific), 500 nM forward and reverse primers (TAG Copenhagen A/S), undiluted cDNA for 5-HT_2B_R, and 10× diluted cDNA for hypoxanthine phosphoribosyl-transferase (Hprt), utilized as a reference gene. Primer sequences were as follows: (mouse) mHPRT-1 Forward 5’-GTTAAGCAGTACAGCCCCAAAATG-3’, Reverse 5’-AAATCCAACAAAGTCTGGCCTGTA-3’; (human) hHPRT-1 Forward 5’ CCCTGGCGTCGTGATTAGTG-3’, Reverse TCGAGCAAGACGTTCAGTCC-3’; (human) h5-HT_2B_R Forward 5’ CTGGTTGGATTGTTTGTGATG-3’, Reverse 5’-CCCTTTAATAGGGACTGGAATG-3’; (mouse) m5-HT_2B_R Forward 5’-CTGATACTCGCGGTGATAATAC-3’, Reverse 5’-TATAGCGGTCCAGGGAAAT-3’; Relative quantification was performed by calculating differences in Ct values between the 5-HT_2B_R and the reference gene across all samples, followed by normalization to the appropriate control group using the 2^−ΔΔCt^ method. Nuclease-free H2O and genomic DNA instead of cDNA were used to control for contamination.

### 4.5 [^3^H]RS127445 autoradiography

Sagittal mouse brain and coronal mouse heart sections, 20 μm in thickness, were collected at 300 μm intervals on ice-cold Superfrost™ slides using a Leica CM3050S cryostat (Nussloch, Germany). The sections were air-dried overnight at 4 °C within a silica gel-containing box and stored at -80 °C until further use.

The 5-HT_2B_-selective ligand [^3^H]RS127445 [56] was used to assess the density of 5-HT_2B_R by real-time autoradiography. Upon thawing at room temperature (RT), the sections were washed in 50-mM Tris-HCl buffer (pH 7.4), containing 150 mM NaCl, 5 mM KCl 1.5 mM MgCl_2_ and 1.5 mM CaCl_2_ (3 × 15 min). Subsequently, the sections were incubated for 3 h in the same buffer, containing 1 nM [^3^H]RS127445 (specific activity 98 Ci/mmol; NT1137 Novandi Chemistry AB). Non-specific binding (NSB) was determined by radiolabelling adjacent sections with 1 nM [^3^H]RS127445 in the presence of 1 μM LY266097 hydrochloride (4081; Tocris). Incubations were terminated by three 1-min washes in ice-cold 50-mM Tris-HCL buffer (pH 7.4), followed by a swift rinse in ice-cold deionized H_2_O (dH2O, Ultra-Clear; Siemens). Sections were then dried under a cold stream of air for 2 h and stored at RT overnight. Autoradiography acquisition was performed for ∼15 h, by using the digital real-time autoradiography system BeaQuant (ai4r, France). Autoradiograms were analysed in a blinded manner by using Beamage software (version 2.1.8, ai4r, France), and a pixel size of 100 μm. The cortex and the hippocampus were manually delineated in 2–6 sections spaced 300 μm apart from each mouse, by reference to the Allen Mouse Brain Atlas for sagittal sections [57]. The gray and white matter of each human neocortical sample was delineated by reference to adjacent, toluidine-blue stained sections. To calculate the mean specific binding (cp/min/mm^2^) of each biological sample, the mean NSB was subtracted from the mean total binding values of all sections analysed.

### 4.6 Statistics

Outliers were identified and removed using the ROUT method. The effects of age and genotype on the specific binding levels of [^3^H]RS127445 and the expression levels of 5-HT_2B_ receptor mRNA were investigated in ageing *APP*_*swe*_*/PS1*_*dE9*_ TG and WT control mice by two-way ANOVA. When significant effects were observed, between-group differences were further explored using Bonferroni multiple comparison *post hoc* tests. Unpaired, two-tailed Student *t*-test was employed to compare levels of 5-HT_2B_R mRNA in AD compared to control samples. Results are presented as mean ± SEM and considered significant when *P*<0.05. All analyses were conducted using GraphPad Prism software (ver. 8.2).

## Supplementary Materials

Figure S1: Digital real-time autoradiography of 5-HT_2B_R binding sites in the cerebellum.

## Author Contributions

Conceptualization, Athanasios Metaxas and Bente Finsen; Formal analysis, Marco Anzalone, Sarmad Karam and Athanasios Metaxas; Funding acquisition, Athanasios Metaxas and Bente Finsen; Investigation, Marco Anzalone, Sarmad Karam, Sanne Briting, Sussanne Petersen, Majken Thomsen and Athanasios Metaxas; Methodology, Marco Anzalone, Sanne Briting, Sussanne Petersen, Majken Thomsen, Anne Landau, Athanasios Metaxas and Bente Finsen; Project administration, Majken Thomsen, Anne Landau and Bente Finsen; Supervision, Athanasios Metaxas and Bente Finsen; Writing – original draft, Marco Anzalone; Writing – review & editing, Sarmad Karam, Sanne Briting, Majken Thomsen, Anne Landau, Athanasios Metaxas and Bente Finsen.

## Institutional Review Board Statement

All procedures complied with Danish law (Bekendtgørelse af lov om dyreforsøg, LBK nr 1107 af 01/07/2022) and European Union directive 2010/63/EU, regulating animal research. Ethical permission was granted by the Animal Ethics Inspectorate of Denmark (nr 2011/561-1950, obtained on 01/04/2011 and 2021-15-0201-00977, obtained on 25/10/2021). Ethical approval for the use of human material was granted by the Danish Biomedical Research Ethical Committee for the Region of Southern Denmark (Project-ID. S-20160036, obtained on 07/07/2016).

## Informed Consent Statement

Written consent forms were obtained for all donors of human tissue.

## Acknowledgments

We acknowledge laboratory technician Janne Skalshøi for the processing of the human brain tissue.

## SUPPLEMENTARY FIGURES

**Supplementary Fig. S1.**
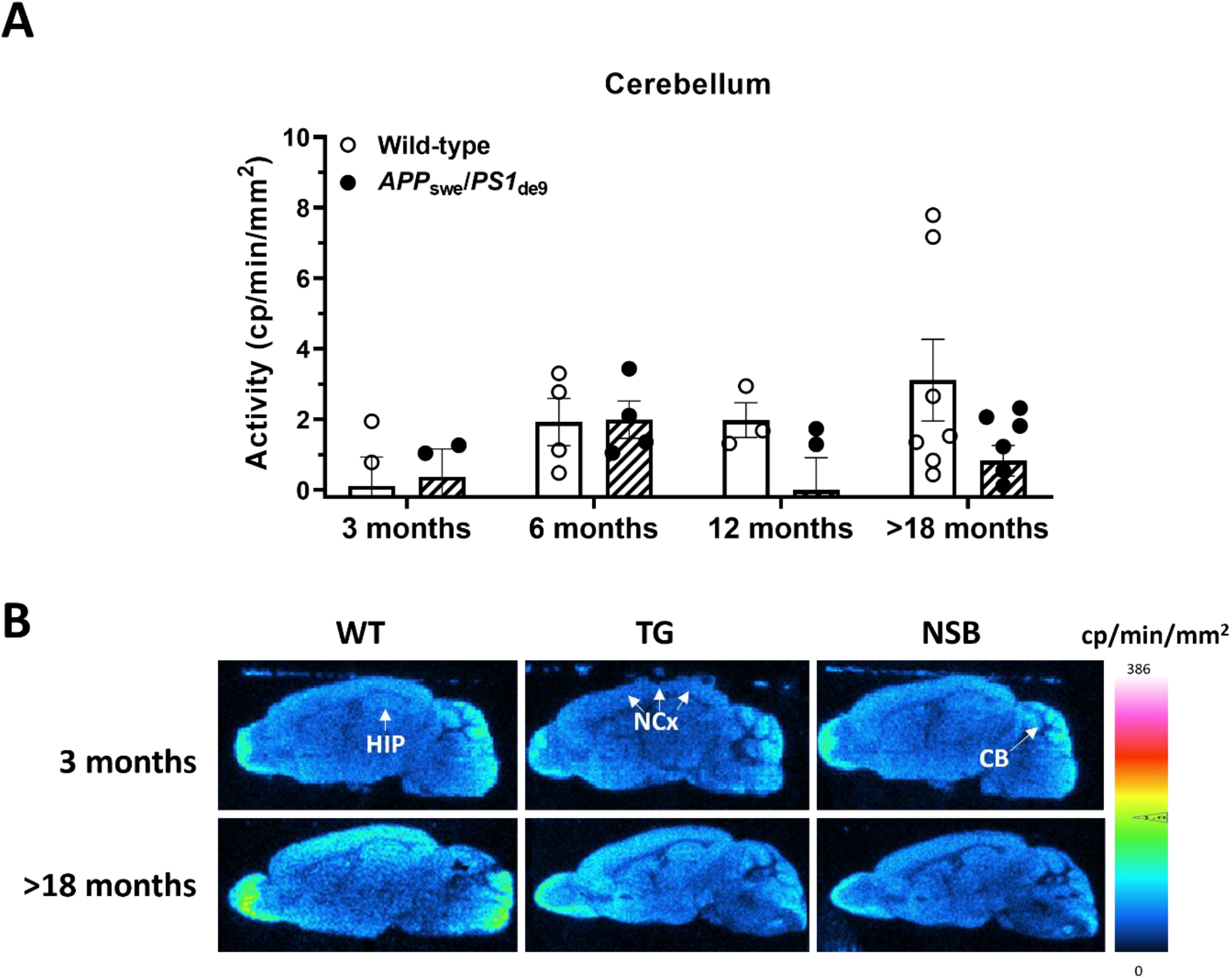
Digital real-time autoradiography of 5-HT_2B_R binding sites in the cerebellum. **(A)** Quantification of [^3^H]RS127445 binding in the cerebellum of ageing WT and *APP*_swe_*/PS1*_dE9_ TG mice. Values represent the mean specific binding of [^3^H]RS127445 ± SEM in 3-8 animals/group, determined in five-six sections from each animal. **(B)** Representative autoradiograms of 5-HT_2B_R binding sites. In 3-month-old mice, specific binding was detected in the cerebellum, but not the neocortex or the hippocampus. Abbreviations: NCx: Neocortex; HIP: Hippocampus; CB: Cerebellum.

## References

[1] M. Foguet et al., ‘Cloning and functional characterization of the rat stomach fundus serotonin receptor.,’ EMBO J., vol. 11, no. 9, p. 3481, Sep. 1992, doi: 10.1002/J.1460-2075.1992.TB05427.X.

[2] J. D. Kursar, D. L. Nelson, D. B. Wainscott, M. L. Cohen, and M. Baez, ‘Molecular cloning, functional expression, and pharmacological characterization of a novel serotonin receptor (5-hydroxytryptamine2F) from rat stomach fundus.,’ Mol. Pharmacol., vol. 42, no. 4, 1992.

[3] K. Schmuck, C. Ullmer, P. Engels, and H. Lübbert, ‘Cloning and functional characterization of the human 5-HT2B serotonin receptor,’ FEBS Lett., vol. 342, no. 1, pp. 85–90, Mar. 1994, doi: 10.1016/0014-5793(94)80590-3.

[4] I. Moutkine, E. L. Collins, C. Béchade, and L. Maroteaux, ‘Evolutionary considerations on 5-HT2 receptors,’ Pharmacol. Res., vol. 140, pp. 14–20, 2019, doi: 10.1016/j.phrs.2018.09.014ï.

[5] P. Bonaventure et al., ‘Nuclei and subnuclei gene expression profiling in mammalian brain,’ Brain Res., vol. 943, no. 1, pp. 38–47, Jul. 2002, doi: 10.1016/S0006-8993(02)02504-0.

[6] D. W. Bonhaus et al., ‘The pharmacology and distribution of human 5-hydroxytryptamine2B (5-HT2B) receptor gene products: comparison with 5-HT2A and 5-HT2c receptors’.

[7] J. D. Kursar, D. L. Nelson, D. B. Wainscott, and M. Baez, ‘Molecular cloning, functional expression, and mRNA tissue distribution of the human 5-hydroxytryptamine2B receptor.,’ Mol. Pharmacol., vol. 46, no. 2, 1994.

[8] D. B. Wainscott, V. L. Lucaites, J. D. Kursar, M. Baez, and D. L. Nelson, ‘Pharmacologic characterization of the human 5-hydroxytryptamine2B receptor: evidence for species differences.,’ J. Pharmacol. Exp. Ther., vol. 276, no. 2, 1996.

[9] S. Loric, J. M. Launay, J. F. Colas, and L. Maroteaux, ‘New mouse 5-HT2-like receptor Expression in brain, heart and intestine,’ FEBS Lett., vol. 312, no. 2–3, pp. 203–207, Nov. 1992, doi: 10.1016/0014-5793(92)80936-B.

[10] M. S. Duxon, T. P. Flanigan, A. C. Reavley, G. S. Baxter, T. P. Blackburn, and K. C. F. Fone, ‘Evidence for expression of the 5-hydroxytryptamine-2B receptor protein in the rat central nervous system,’ Neuroscience, vol. 76, no. 2, pp. 323–329, 1997, doi: 10.1016/S0306-4522(96)00480-0.

[11] T. F. Galatro et al., ‘Transcriptomic analysis of purified human cortical microglia reveals age-associated changes,’ Nat. Neurosci., vol. 20, no. 8, pp. 1162–1171, Aug. 2017, doi: 10.1038/NN.4597.

[12] D. Gosselin et al., ‘An environment-dependent transcriptional network specifies human microglia identity,’ Science, vol. 356, no. 6344, pp. 1248–1259, Jun. 2017, doi: 10.1126/SCIENCE.AAL3222.

[13] M. Benekareddy, K. C. Vadodaria, A. R. Nair, and V. A. Vaidya, ‘Postnatal serotonin type 2 receptor blockade prevents the emergence of anxiety behavior, dysregulated stress-induced immediate early gene responses, and specific transcriptional changes that arise following early life stress,’ Biol. Psychiatry, vol. 70, no. 11, pp. 1024–1032, Dec. 2011, doi: 10.1016/J.BIOPSYCH.2011.08.005.

[14] R. Tikkanen et al., ‘Impulsive alcohol-related risk-behavior and emotional dysregulation among individuals with a serotonin 2B receptor stop codon,’ Transl. Psychiatry 2015 511, vol. 5, no. 11, pp. e681–e681, Nov. 2015, doi: 10.1038/tp.2015.170.

[15] P. M. Pitychoutis, A. Belmer, I. Moutkine, J. Adrien, and L. Maroteaux, ‘Mice Lacking the Serotonin Htr2B Receptor Gene Present an Antipsychotic-Sensitive Schizophrenic-Like Phenotype,’ Neuropsychopharmacology, vol. 40, no. 12, pp. 2764–2773, Nov. 2015, doi: 10.1038/NPP.2015.126.

[16] D. Goldman et al., ‘A population-specific HTR2B stop codon predisposes to severe impulsivity,’ Nature, vol. 468, no. 7327, pp. 1061–1068, Dec. 2010, doi: 10.1038/NATURE09629.

[17] S. Chakraborty, J. C. Lennon, S. A. Malkaram, Y. Zeng, D. W. Fisher, and H. Dong, ‘Serotonergic System, Cognition, and BPSD in Alzheimer’s disease’, doi: 10.1016/j.neulet.2019.03.050.

[18] C. U. Von Linstow et al., ‘Effect of aging and Alzheimer’s disease-like pathology on brain monoamines in mice,’ Neurochem. Int., vol. 108, pp. 238–245, Sep. 2017, doi: 10.1016/J.NEUINT.2017.04.008.

[19] G. S. Smith et al., ‘Molecular imaging of serotonin degeneration in mild cognitive impairment,’ Neurobiol. Dis., vol. 105, pp. 33–41, 2017, doi: 10.1016/j.nbd.2017.05.007.

[20] S. G. Snowden et al., ‘Neurotransmitter Imbalance in the Brain and Alzheimer’s Disease Pathology,’ J. Alzheimers. Dis., vol. 72, no. 1, pp. 35–43, 2019, doi: 10.3233/JAD-190577.

[21] H. Aral, K. Kosaka, and R. Iizuka, ‘Changes of biogenic amines and their metabolites in postmortem brains from patients with Alzheimer-type dementia,’ J. Neurochem., vol. 43, no. 2, pp. 388–393, 1984, doi: 10.1111/J.1471-4159.1984.TB00913.X.

[22] M. Hendricksen, A. J. Thomas, I. N. Ferrier, P. Ince, and J. T. O’Brien, ‘Neuropathological study of the dorsal raphe nuclei in late-life depression and Alzheimer’s disease with and without depression,’ Am. J. Psychiatry, 2004, doi: 10.1176/appi.ajp.161.6.1096.

[23] D. Š. Štrac, N. Pivac, and D. Mück-Šeler, ‘The serotonergic system and cognitive function,’ Transl. Neurosci., vol. 7, no. 1, p. 35, Jan. 2016, doi: 10.1515/TNSCI-2016-0007.

[24] P. Holm et al., ‘Plaque deposition dependent decrease in 5-HT2A serotonin receptor in AbetaPPswe/PS1dE9 amyloid overexpressing mice,’ J. Alzheimers. Dis., vol. 20, no. 4, pp. 1201–1213, 2010, doi: 10.3233/JAD-2010-100117.

[25] M. K. P. Lai et al., ‘Reduced serotonin 5-HT1A receptor binding in the temporal cortex correlates with aggressive behavior in Alzheimer disease,’ Brain Res., vol. 974, no. 1–2, pp. 82–87, Jun. 2003, doi: 10.1016/S0006-8993(03)02554-X.

[26] D. M. Bowen, A. Najlerahim, A. W. Procter, P. T. Francis, and E. Murphy, ‘Circumscribed changes of the cerebral cortex in neuropsychiatric disorders of later life,’ Proc. Natl. Acad. Sci. U. S. A., vol. 86, no. 23, pp. 9504–9508, 1989, doi: 10.1073/PNAS.86.23.9504.

[27] A. V. T. Cheng et al., ‘Cortical serotonin-S2 receptor binding in Lewy body dementia, Alzheimer’s and Parkinson’s diseases,’ J. Neurol. Sci., vol. 106, no. 1, pp. 50–55, 1991, doi: 10.1016/0022-510X(91)90193-B.

[28] M. K. Lai et al., ‘Loss of serotonin 5-HT2A receptors in the postmortem temporal cortex correlates with rate of cognitive decline in Alzheimer’s disease,’ Psychopharmacology (Berl)., vol. 179, no. 3, pp. 673–677, May 2005, doi: 10.1007/S00213-004-2077-2/METRICS.

[29] D. E. Lorke, G. Lu, E. Cho, and D. T. Yew, ‘Serotonin 5-HT2A and 5-HT6 receptors in the prefrontal cortex of Alzheimer and normal aging patients,’ BMC Neurosci., vol. 7, no. 1, pp. 1–8, Apr. 2006, doi: 10.1186/1471-2202-7-36/FIGURES/3.

[30] C. P. L. H. Chen et al., ‘Presynaptic Serotonergic Markers in Community-Acquired Cases of Alzheimer’s Disease: Correlations with Depression and Neuroleptic Medication,’ J. Neurochem., vol. 66, no. 4, pp. 1592–1598, Apr. 1996, doi: 10.1046/J.1471-4159.1996.66041592.X.

[31] A. Metaxas, M. Anzalone, R. Vaitheeswaran, S. Petersen, A. M. Landau, and B. Finsen, ‘Neuroinflammation and amyloid-beta 40 are associated with reduced serotonin transporter (SERT) activity in a transgenic model of familial Alzheimer’s disease,’ Alzheimers. Res. Ther., vol. 11, no. 1, May 2019, doi: 10.1186/S13195-019-0491-2.

[32] G. Mengod, H. Nguyen, H. Le, C. Waeber, H. Lübbert, and J. M. Palacios, ‘The distribution and cellular localization of the serotonin 1C receptor mRNA in the rodent brain examined by in situ hybridization histochemistry. Comparison with receptor binding distribution,’ Neuroscience, vol. 35, no. 3, pp. 577–591, 1990, doi: 10.1016/0306-4522(90)90330-7.

[33] D. S. Choi and L. Maroteaux, ‘Immunohistochemical localisation of the serotonin 5-HT2B receptor in mouse gut, cardiovascular system, and brain,’ FEBS Lett., vol. 391, no. 1–2, pp. 45–51, Aug. 1996, doi: 10.1016/0014-5793(96)00695-3.

[34] W. Wisden et al., ‘Cloning and characterization of the rat 5-HT5B receptor. Evidence that the 5-HT5B receptor couples to a G protein in mammalian cell membranes,’ FEBS Lett., vol. 333, no. 1–2, pp. 25–31, Oct. 1993, doi: 10.1016/0014-5793(93)80368-5.

[35] M. Pompeiano, J. M. Palacios, and G. Mengod, ‘Distribution of the serotonin 5-HT2 receptor family mRNAs: comparison between 5-HT2A and 5-HT2C receptors,’ Mol. Brain Res., vol. 23, no. 1–2, pp. 163–178, Apr. 1994, doi: 10.1016/0169-328X(94)90223-2.

[36] M. Baez, J. D. Kursar, L. A. Helton, D. B. Wainscott, and D. L. Nelson, ‘Molecular biology of serotonin receptors.,’ Obes. Res., vol. 3 Suppl 4, 1995, doi: 10.1002/J.1550-8528.1995.TB00211.X.

[37] L. A. Helton, K. B. Thor, and M. Baez, ‘5-Hydroxytryptamine2A, 5-hydroxytryptamine2B, and 5-hydroxytryptamine2C receptor mRNA expression in the spinal cord of rat, cat, monkey and human,’ Neuroreport, vol. 5, no. 18. pp. 2617–2620, Dec. 01, 1994. doi: 10.1097%2F00001756-199412000-00053.

[38] J. F. López-Giménez, L. H. Tecott, J. M. Palacios, G. Mengod, and M. T. Vilaró, ‘Serotonin 5-HT (2C) receptor knockout mice: autoradiographic analysis of multiple serotonin receptors,’ J. Neurosci. Res., vol. 67, no. 1, pp. 69–85, Jan. 2002, doi: 10.1002/JNR.10072.

[39] D. W. Bonhaus et al., ‘The pharmacology and distribution of human 5-hydroxytryptamine2B (5-HT2b) receptor gene products: comparison with 5-HT2a and 5-HT2c receptors,’ Br. J. Pharmacol., vol. 115, no. 4, pp. 622–628, 1995, doi: 10.1111/J.1476-5381.1995.TB14977.X.

[40] P. Bonaventure et al., ‘Molecular and pharmacological characterization of serotonin 5-HT 2A and 5-HT2B receptor subtypes in dog,’ Eur. J. Pharmacol., vol. 513, no. 3, pp. 181–192, Apr. 2005, doi: 10.1016/j.ejphar.2005.03.013.

[41] S. F. Tadros, M. D’Souza, M. L. Zettel, X. X. Zhu, M. Lynch-Erhardt, and R. D. Frisina, ‘Serotonin 2B receptor: Upregulated with age and hearing loss in mouse auditory system,’ Neurobiol. Aging, vol. 28, no. 7, pp. 1112–1123, Jul. 2007, doi: 10.1016/J.NEUROBIOLAGING.2006.05.021.

[42] V. G. Hernandez et al., ‘Translatome analysis reveals microglia and astrocytes to be distinct regulators of inflammation in the hyperacute and acute phases after stroke,’ Glia, vol. 71, no. 8, pp. 1960–1984, Aug. 2023, doi: 10.1002/GLIA.24377.

[43] Y. Zhang et al., ‘An RNA-Sequencing Transcriptome and Splicing Database of Glia, Neurons, and Vascular Cells of the Cerebral Cortex,’ J. Neurosci., vol. 34, no. 36, pp. 11929–11947, Sep. 2014, doi: 10.1523/JNEUROSCI.1860-14.2014.

[44] L. Maroteaux and L. Monassier, Eds., ‘5-HT2B Receptors,’ vol. 35, 2021, doi: 10.1007/978-3-030-55920-5.

[45] Y. Zhang et al., ‘Purification and Characterization of Progenitor and Mature Human Astrocytes Reveals Transcriptional and Functional Differences with Mouse,’ Neuron, vol. 89, no. 1, pp. 37–53, Jan. 2016, doi: 10.1016/j.neuron.2015.11.013.

[46] Y. Liu et al., ‘Amyloid Pathology Is Associated with Progressive Monoaminergic Neurodegeneration in a Transgenic Mouse Model of Alzheimer’s Disease,’ J. Neurosci., vol. 28, no. 51, p. 13805, Dec. 2008, doi: 10.1523/JNEUROSCI.4218-08.2008.

[47] G. Krabbe, V. Matyash, U. Pannasch, L. Mamer, H. W. G. M. Boddeke, and H. Kettenmann, ‘Activation of serotonin receptors promotes microglial injury-induced motility but attenuates phagocytic activity,’ Brain. Behav. Immun., vol. 26, no. 3, pp. 419–428, Mar. 2012, doi: 10.1016/J.BBI.2011.12.002.

[48] A. A. Babcock et al., ‘Cytokine-producing microglia have an altered beta-amyloid load in aged APP/PS1 Tg mice,’ Brain. Behav. Immun., vol. 48, pp. 86–101, 2015, doi: 10.1016/J.BBI.2015.03.006.

[49] C. L. Myhre et al., ‘Microglia Express Insulin-Like Growth Factor-1 in the Hippocampus of Aged APPswe/PS1ΔE9 Transgenic Mice,’ Front. Cell. Neurosci., vol. 13, p. 438590, Jul. 2019, doi: 10.3389/FNCEL.2019.00308/BIBTEX.

[50] C. Béchade et al., ‘The serotonin 2B receptor is required in neonatal microglia to limit neuroinflammation and sickness behavior in adulthood,’ Glia, vol. 69, no. 3, pp. 638–654, Mar. 2021, doi: 10.1002/GLIA.23918.

[51] A. M. Buga et al., ‘Up-regulation of serotonin receptor 2B mRNA and protein in the peri-infarcted area of aged rats and stroke patients,’ Oncotarget, vol. 7, no. 14, pp. 17415–17430, 2016, doi: 10.18632/ONCOTARGET.8277.

[52] A. Cathala, C. Devroye, G. Drutel, J. M. Revest, F. Artigas, and U. Spampinato, ‘Serotonin2B receptors in the rat dorsal raphe nucleus exert a GABA-mediated tonic inhibitory control on serotonin neurons,’ Exp. Neurol., vol. 311, pp. 57–66, Jan. 2019, doi: 10.1016/J.EXPNEUROL.2018.09.015.

[53] A. L. Auclair et al., ‘The central serotonin 2B receptor: a new pharmacological target to modulate the mesoaccumbens dopaminergic pathway activity,’ J. Neurochem., vol. 114, no. 5, pp. 1323–1332, Sep. 2010, doi: 10.1111/J.1471-4159.2010.06848.X.

[54] A. Benhadda et al., ‘Serotonin 2B Receptor by Interacting with NMDA Receptor and CIPP Protein Complex May Control Structural Plasticity at Glutamatergic Synapses,’ ACS Chem. Neurosci., vol. 12, no. 7, pp. 1133–1149, Apr. 2021, doi: 10.1021/ACSCHEMNEURO.0C00638.

[55] J. L. Jankowsky, H. H. Slunt, V. Gonzales, N. A. Jenkins, N. G. Copeland, and D. R. Borchelt, ‘APP processing and amyloid deposition in mice haplo-insufficient for presenilin 1,’ Neurobiol. Aging, vol. 25, no. 7, pp. 885–892, Aug. 2004, doi: 10.1016/j.neurobiolaging.2003.09.008.

[56] D. W. Bonhaus et al., ‘RS-127445: a selective, high affinity, orally bioavailable 5-HT2B receptor antagonist,’ Br. J. Pharmacol., vol. 127, no. 5, p. 1075, 1999, doi: 10.1038/SJ.BJP.0702632.

[57] ‘Reference Atlas:: Allen Brain Atlas: Mouse Brain.’ Accessed: Dec. 11, 2023. [Online]. Available: https://mouse.brain-map.org/static/atlas

